# Inference of cell-type specific imprinted regulatory elements and genes during human neuronal differentiation

**DOI:** 10.1101/2021.10.04.463060

**Authors:** Dan Liang, Nil Aygün, Nana Matoba, Folami Y. Ideraabdullah, Michael I. Love, Jason L. Stein

## Abstract

Genomic imprinting results in gene expression biased by parental chromosome of origin and occurs in genes with important roles during human brain development. However, the cell-type and temporal specificity of imprinting during human neurogenesis is generally unknown. By detecting within-donor allelic biases in chromatin accessibility and gene expression that are unrelated to cross-donor genotype, we inferred imprinting in both primary human neural progenitor cells (phNPCs) and their differentiated neuronal progeny from up to 85 donors. We identified 43/20 putatively imprinted regulatory elements (IREs) in neurons/progenitors, and 133/79 putatively imprinted genes in neurons/progenitors. Though 10 IREs and 42 genes were shared between neurons and progenitors, most imprinting was only detected within specific cell types. In addition to well-known imprinted genes and their promoters, we inferred novel IREs and imprinted genes. We found IREs overlapped with CpG islands more than non-imprinted regulatory elements. Consistent with DNA methylation-based regulation of imprinted expression, some putatively imprinted regulatory elements also overlapped with differentially methylated regions on the maternal germline. Finally, we identified a progenitor-specific putatively imprinted gene overlap with copy number variation that is associated with uniparental disomy-like phenotypes. Our results can therefore be useful in interpreting the function of variants identified in future parent-of-origin association studies.

## INTRODUCTION

In contrast to most loci in the genome that have roughly equal expression from either parental chromosome, genomic imprinting leads to biased levels of gene expression or chromatin accessibility from either the maternal or paternal chromosome. Some genomic imprinting results in totally silenced expression of one parental allele and is more likely to be shared across multiple tissues (Bonthuis et al. 2015). However, most imprinted genes exhibit some tissue and cell-type specific imprinted expression (Bonthuis et al. 2015; Kravitz and Gregg 2019; Zink et al. 2018). Some genes are imprinted specifically in humans (Nakabayashi et al. 2011) and many imprinted genes are also expressed in neural development or in the adult brain (Babak et al. 2015; Barlow and Bartolomei 2014; Perez, Rubinstein, and Dulac 2016). Previous studies have identified imprinted genes in different human tissues or cells (Babak et al. 2015; Baran et al. 2015; Santoni et al. 2017), but imprinted regulatory elements (IREs) have not been well defined during human neurogenesis nor has their cell-type specificity been assessed. Elucidation of cell-type specific imprinting mechanisms during neurogenesis are critical for interpreting results of parent-of-origin association studies for neuropsychiatric disorders and subsequent therapeutic development (Perez, Rubinstein, and Dulac 2016; Ishida and Moore 2013; Nicholls 2000; Mozaffari et al. 2019; Brandler et al. 2018; Wolter et al. 2020).

The primary regulatory elements (REs) that control genomic imprinting, called imprinting control elements (ICEs) or imprinting control regions (ICRs), exhibit parental specific DNA methylation. Some of these differentially methylated regions (DMRs) are inherited from the sperm or egg and are maintained postfertilization throughout development in all tissues (Ferguson-Smith 2011; Hanna and Kelsey 2014; Arnaud 2010; Plasschaert and Bartolomei 2014). A separate class of DMRs acquire methylation after fertilization under the direction of germ-cell specific DMRs and show tissue-specific methylation patterns (Lopes et al. 2003; Lucifero et al. 2002). Cell-type specific “reading” of germline DMRs and other epigenetic alterations such as histone modifications without associated germ-cell specific DMRs also contribute to regulation of imprinted expression in specific tissues/cell-types (Prickett and Oakey 2012; Arnaud 2010; Andergassen et al. 2017; Xu Wang, Soloway, and Clark 2011; Q. Wang et al. 2011). However, these tissue/cell-type specific regulatory elements controlling imprinting are not well identified.

Genomic imprinting is most often studied in mice where reciprocal cross breeding is designed, parents are genotyped, and parent-of-origin specific gene expression is measured in offspring using high-throughput sequencing data (X. Wang and Clark 2014; Oreper et al. 2018; Crowley et al. 2015). Heterozygous genetic markers then allow the separation of maternal vs paternal expression (Babak et al. 2008; Xu Wang et al. 2008; Gregg et al. 2010; Laukoter et al. 2020). These studies have found that there are more genes with imprinted expression in the brain as compared to non-brain tissues (Perez et al. 2015) and that genomic imprinting shows key functions in neurodevelopmental processes including neural progenitor expansion, migration, differentiation, and cell polarization in mouse brains (Perez, Rubinstein, and Dulac 2016). However, in humans, it is difficult to obtain both molecular phenotypes and genotype data from children and genotype data from both parents on a large scale in order to demonstrate genomic imprinting. It is still possible to infer imprinting expression in humans where parental genotypes are unavailable by detecting within donor allelic imbalance unrelated to across donor genotype in large datasets where sequencing data for gene expression/chromatin accessibility and genotyping have been collected (Figure 1A) (Reinius and Sandberg 2015; Baran et al. 2015). Here, we used the beta-binomial distribution to model allelic counts across a population up to 85 genetically diverse donors, in order to estimate the dispersion of the allelic ratio (AR; reference allelic count to total count) in the population, for gene expression (RNA-seq) and chromatin accessibility (ATAC-seq). Higher dispersion of the allelic ratio is suggestive of parentally biased expression or accessibility as the parental allele is not expected to be consistent with reference allele coding. Using this method, we identify putatively IREs and genes in human neural progenitors and neurons.

**Figure 1.**
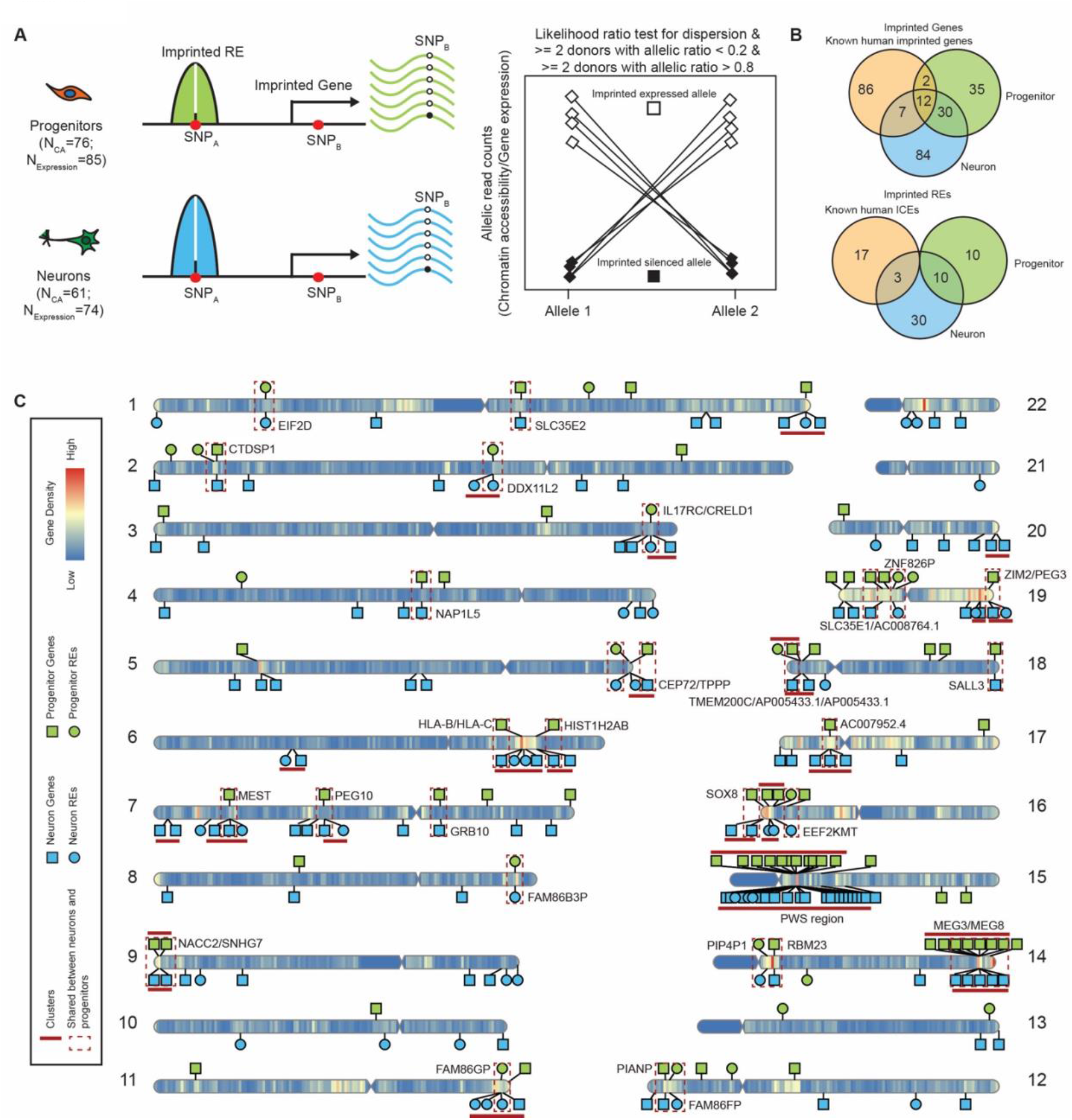
Identification of neuron/progenitor imprinted genes and Res. (A)Schematic cartoon of experimental design and methods.(B) Comparison of imprinted genes and REs in neurons, progenitors and known imprinted genes/ICEs.(C) Ideogram of neuron/progenitor imprinted genes and REs on the human genome.

## RESULTS

### Inference of IREs and genes in progenitors and neurons

We utilized primary human neural progenitor (phNPC) chromatin accessibility and expression quantitative trait loci (QTL) datasets described in our previous work (Liang et al. 2021; Aygün et al. 2021) to infer imprinted genes and REs. phNPCs were cultured *in vitro* as progenitor cells and also differentiated for 8 weeks, virally labeled, and sorted to obtain a homogeneous population of neurons. We then performed ATAC-seq and RNA-seq to obtain chromatin accessibility profiles (Liang et al. 2021) (N_Progenitor=76 and N_Neuron=61) and gene expression profiles (Aygün et al. 2021) (N_Progenitor=85 and N_Neuron=74) in both cell types (Figure 1A). We identified heterozygous genetic variants using imputed genotype data for all donors in progenitors and neurons in order to distinguish the two chromosomes, though we are unable to classify maternal or paternal origin. Allele-specific chromatin accessibility and gene expression were then calculated using read counts (for the reference allele and the alternative allele) at each accessible/expressed heterozygous single nucleotide polymorphism (SNP) site. We identified 19,960 heterozygous SNPs in 12,233 accessible regions and 80,811 expressed heterozygous SNPs in 11,770 genes in neurons; and 42,706 heterozygous SNPs in 10,802 accessible regions and 63,733 expressed heterozygous SNPs in 8,958 genes in progenitors.

The silenced imprinted allele, inherited from one parent, will by definition lead to decreased chromatin accessibility/expression as compared to the imprinted expressed allele. Here, we do not know the parental origin of each allele at a heterozygous locus. We expect that at an imprinted site, the AR will be either high (when the reference allele is the expressed imprinted allele) or low (when the reference allele is the silenced imprinted allele) leading to high dispersion in estimates of the AR across donors. We estimated the dispersion of allelically biased chromatin accessibility or gene expression to identify the SNPs with highly variable (either high or low) AR in the population. We modeled allelic counts of chromatin accessibility or gene expression using a beta-binomial distribution at heterozygous SNPs (Skelly et al. 2011; Castel et al. 2015) and evaluated the significance of the over-dispersion of AR using a likelihood ratio test. SNPs in known randomly monoallelic expression (RMAE) genes and randomly monoallelic chromatin accessible (RMACA) regions were removed (Xu et al. 2017; Gimelbrant et al. 2007). Finally, we kept SNPs with a significant AR over-dispersion (FDR < 0.05 across all heterozygous SNPs tested within a given cell type (Benjamini and Hochberg 1995)). A minimal donor count for both high and low AR (n_donor_ ≥ 2 for AR ≥ 0.8 and n_donor_ ≥ 2 for AR ≤ 0.2) was used to exclude cases of allele specific expression or chromatin accessibility, i.e. AR driven by genetic effects (Figure 1A). Because our study lacks parental genotype data, we cannot exclude the possibility that allelic bias in newly identified imprinted genes/REs is due to previously unidentified RMAE or RMACA rather than parental inheritance. Thus, we refer to these allele-specifically expressed genes/REs as putatively imprinted.

After these analyses and filtering steps, we identified 43 IREs containing 57 SNPs in neurons (nIREs) and 20 IREs containing 25 SNPs in progenitors (pIREs) (Figure 1B-1C; Supplementary Figure 1A, 1C and 1D; Supplemental Table S1). We found 3 nIREs overlapped with known human ICEs for known imprinted genes *PEG10, MEST* and *ZIM2/PEG3 (Cowley et al. 2018)*, providing confidence in IRE calls. We also found 10 shared nIREs and pIREs overlapped with the promoters of 11 well-known imprinted genes (Supplemental Figure 1A; red labeled points) that are involved in neuronal development and differentiation: *MAGEL2, NDN, SGCE, PEG10, NAA60, MIMT1, ZNF597, MEST, PEG3, ZIM2* and *SNRPN* (Watrin et al. 2005; Grütz et al. 2017; Ono et al. 2006; Babak et al. 2015; Nakabayashi et al. 2011). Surprisingly, we did not find any pIREs that overlapped with the promoters of well-known imprinted genes, so the pIREs identified here may represent a novel set of imprinted elements, such as the promoters of *EIF2D, DDX11L2* and *SPEG*.

We identified 133 neuron imprinted genes (20 of which are previously known imprinted genes) containing 653 SNPs and 79 progenitor imprinted genes (15 of which are previously known imprinted genes) containing 166 SNPs (Figure 1B-1C; Supplemental Figure 1B; Supplemental Table S2). For these imprinted genes, many genes have been previously reported as imprinted, such as *UBE3A, NDN, PEG3, MEST, GRB10* and *MEG3* (Dindot et al. 2008; Huntriss et al. 2013; Blagitko et al. 2000; Zhang et al. 2003; Jay et al. 1997; Ho-Shing and Dulac 2019). Imprinted genes like *DLK1* and *ZDBF2* have previously described important functions in corticogenesis (Duffié et al. 2014; Bouschet et al. 2017; Ferrón et al. 2011; Surmacz et al. 2012). We also found novel imprinted genes specifically in neurons, such as *HM13, ZNF331, COPG2, DOC2B* and *PBX1* (Supplemental Table S2) that have not been previously described as imprinted but where the expression patterns fit the characteristics of imprinting. However, we could not detect all of the known imprinted genes, such as *IGF2*, due to lack of expression in these two cell types (GTEx Consortium 2020; Nowakowski et al. 2017).

We found 42 genes that show imprinting patterns in both progenitors and neurons (Figure 1B, Supplementary Table 1), including the well-known imprinted genes: *GRB10, PEG10, MEST, MEG3, MEG8, NDN*, and *SNHG14*. Several of the imprinted genes that are shared across cell types have not been previously identified as imprinted, such as *PIANP* and *SNHG7*. We identified 10 REs showing evidence of imprinting in both progenitors and neurons, at the promoters of *EIF2D, DDX11L2, IL17RC, CRELD1, FAM86B3P, FAM86GP, FAM86FP, PIP4P1, EEF2KMT* and *ZNF826P*. Many of these REs overlapped with promoters of genes that are not previously known as imprinted genes, such as *CRELD1* and *EIF2D*, which could be new candidates for imprinting regulation. Though there was some overlap of imprinting between cell types (31.6%/53.2% imprinted genes and 23.3%/50.0% IREs in neurons/progenitors), most imprinting was found only in one cell type. Often this was because the gene did not survive QC in both cell types (Supplementary Figure 1C and 1D). Nevertheless, cell type specific imprinting patterns were still detectable when the same gene passed QC in both cell types. Overall, the detection of well-known imprinted genes and regulatory elements supports the statistical approach to identify imprinted candidates outlined here. Novel genes and regulatory elements may have been undetected in previous studies due to the lack of a cell type specific or development system for studying these effects.

### Neuron/progenitor imprinting at known loci

A well-known example of an imprinted genomic cluster is the Prader–Willi/Angelman Syndrome (PWS/AS) region on human chromosome 15q11–q13 (Nicholls, Saitoh, and Horsthemke 1998). Mutations in the PWS/AS region result in neurodevelopmental disorders, PWS and AS (Perk et al. 2002), in a parent-of-origin dependent manner, demonstrating the important function of these imprinted genes during neural development. We identified neuron-specific IREs that overlap with the promoters of *MAGEL2, NDN* and *SNRPN*, the latter with multiple SNPs supporting the imprinting inference (Figure 1C, 2A and 2B). 72 SNPs in *UBE3A* passed our filtering criteria and showed evidence for neuron-specific imprinting of gene expression (Figure 2A and 2C; Supplemental Figure 2A). This finding is in agreement with previous studies showing that *UBE3A* is expressed exclusively from the maternally inherited allele in neurons (Martins-Taylor et al. 2014; Hsiao et al. 2019). In the PWS/AS region, we found more IREs and genes in neurons than in progenitors, which is also consistent with previous studies in iPSC-derived neurons (Pólvora-Brandão et al. 2018; Stanurova et al. 2016).

**Figure 2.**
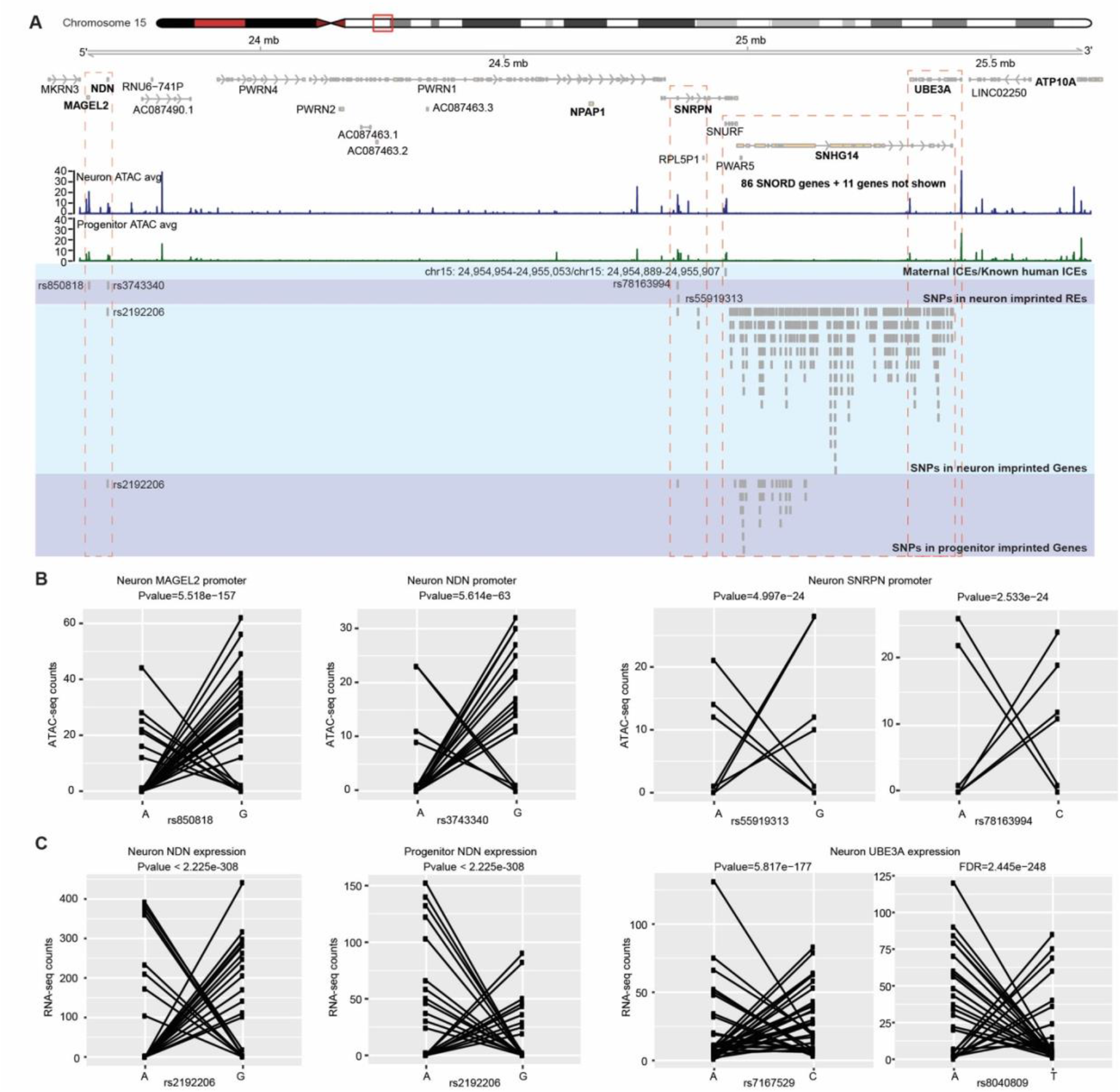
Imprinted genes and REs at known genomic imprinting loci. (A) Coverage plot of ATAC-seq in neurons and progenitors at the AS/PWS lous and SNPs in the imprinted REs and genes. (B) Allelic ATAC-seq counts for selected SNPs in imprinted REs in (A). (C) Allelic RNA-seq counts for selected SNPs in imprinted genes in (A).

### Methylation and transcription factor binding at IREs

The methylation of CpG sites is an important epigenetic regulation of imprinting in mammals (Paulsen and Ferguson-Smith 2001). To explore the relationship between methylation and IREs in neurons and progenitors, we first calculated the GC content of the IREs. We found the GC content is significantly higher within IREs as compared to non-imprinted REs (Figure 3A). We found the IREs showed significantly higher overlap with human CpG islands than the non-imprinted REs in both neurons and progenitors (Figure 3B). Enrichment of CpG sites at IREs supports a role for DNA methylation in genomic imprinting at these loci during neuronal differentiation.

**Figure 3.**
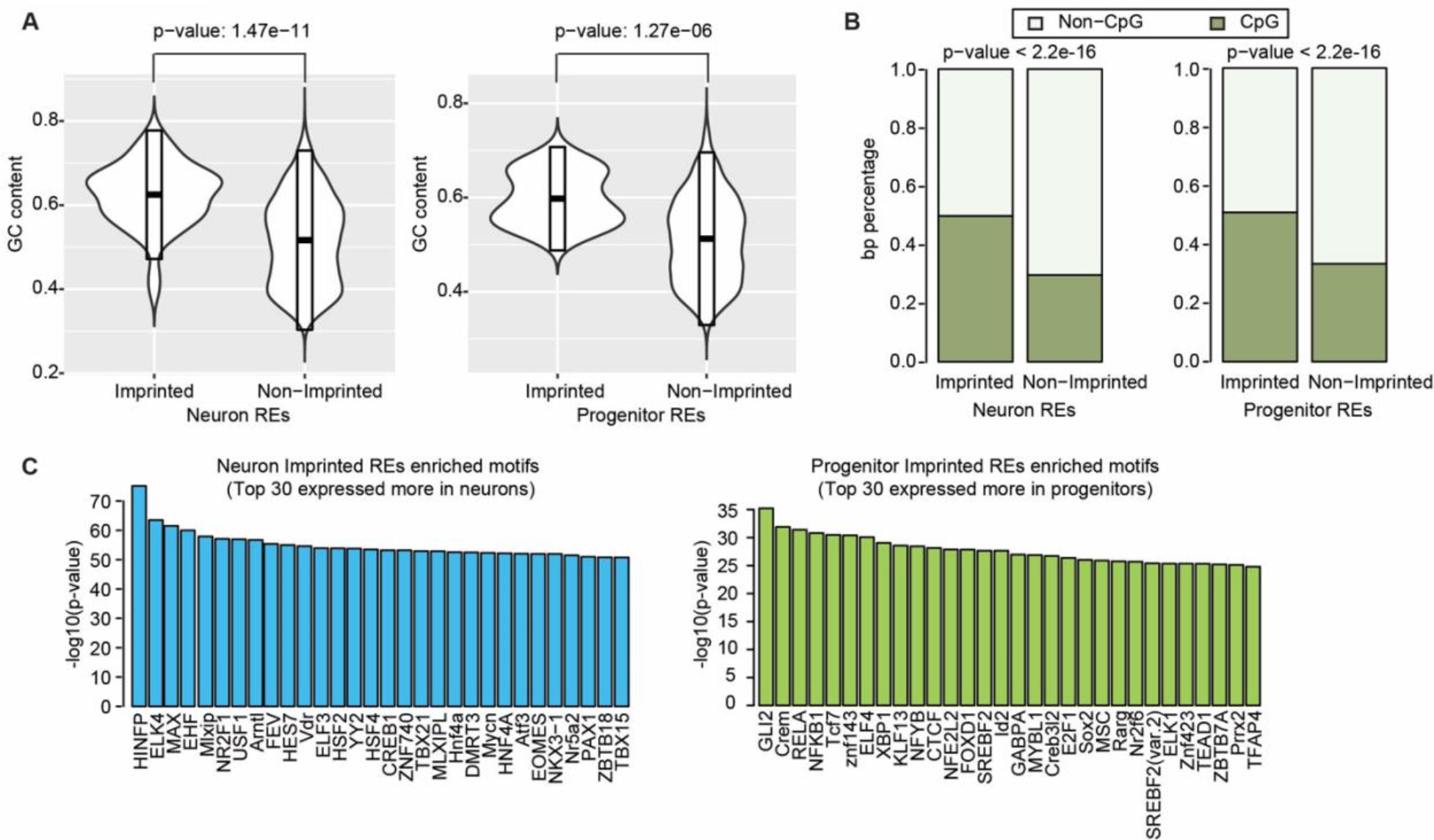
Methylation and transcription factor binding at imprinted REs. (A) GC content of imprinted and non-imprinted REs in neurons (*left*) and progenitors (*right*). (B) Overlap of imprinted and non-imprinted REs with human CpG islands in neurons (*left*) and progenitors (*right*). (C) Enriched TF motifs within imprinted REs as compared to non-imprinted REs.

DNA-binding proteins (DBP) binding to the IREs have important roles in maintaining imprinting by changing the methylation levels around their binding sites during early development (Takahashi et al. 2019; Sanli and Feil 2015). Additionally, methylation at IREs may block some DBPs binding on the imprinted silenced allele thereby decreasing expression of genes from the imprinted silenced allele. However, it is unclear which DBPs are involved in either maintaining imprinting or the downstream consequences of imprinting during human neurogenesis, especially at cell-type resolution. To identify the DBPs that may bind to and regulate IREs in neurons and progenitors, we performed an enrichment analysis of transcription factor (TF) motifs in the IREs using a binomial test (McLean et al. 2010). We retained only the TFs with significantly higher expression in the cell type tested for enrichment (Figure 3C). Among TF motifs enriched in neuron IREs, we found ELK4, which is involved in upstream regulation of parent-of-origin-regulated genes in mice with sleep loss (Tinarelli et al. 2014). We also found that CTCF TF motifs were enriched in pIREs. CTCF was shown to regulate the imprinted expression of *KLD1* by binding to the ICE at *KLD1-MEG3* locus in embryo stem cells (Llères et al. 2019). Using these IREs, we are able to identify TFs implicated in imprinting gene regulation during human neurogenesis.

### Imprinted REs and genes indicate isoform-specific imprinting in neurons

Previous studies showed isoform-specific imprinted transcription for well known genes, like *PEG1* and *MEST* (Kosaki et al. 2000; Stelzer et al. 2015). However, the number of isoform-specific imprinted genes may still be underestimated, because isoform expression is highly cell type specific and imprinting has not been assessed in all cell types. Using IREs in neurons and progenitors, we predicted isoform-specific imprinted expression in each cell type. In neurons, we found that the promoter region (chr21:39,385,651-39,386,540) of one isoform (ENST00000380713) of the gene *GET1* (also known as *WRB*) showed an imprinting pattern at two SNPs (Figure 4A and 4B). We did not find any SNP passed QC in this RE in progenitors, therefore we are not able to test if this RE is a pIRE. This region in *WRB* was previously reported as a new candidate imprinted region according to DNA methylation analysis using peripheral blood samples (Docherty et al. 2014; Alves da Silva et al. 2016), but to our knowledge, this is the first time this region was suggested as an IRE in neurons. WRB is a receptor associated with protein transmembrane transport (Vilardi, Lorenz, and Dobberstein 2011); but, its function in the human brain is unknown. This region overlapped with a differentially methylated region that is more methylated in oocyte and morula than sperm and maintains methylation in fetal and adolescent brain (Figure 4C; Supplementary Figure 2B-C) (Guo et al. 2014; Lister et al. 2013). The DMR inherited from germ cells in the soma provides evidence that this IRE is an ICE that is maintained throughout development and potentially regulates maternal genomic imprinting of a specific *GET1* isoform. No SNPs in the expression level data survived QC so we could not test whether *GET1* isoform expression was imprinted (Plasschaert and Bartolomei 2014; Hanna and Kelsey 2014).

**Figure 4.**
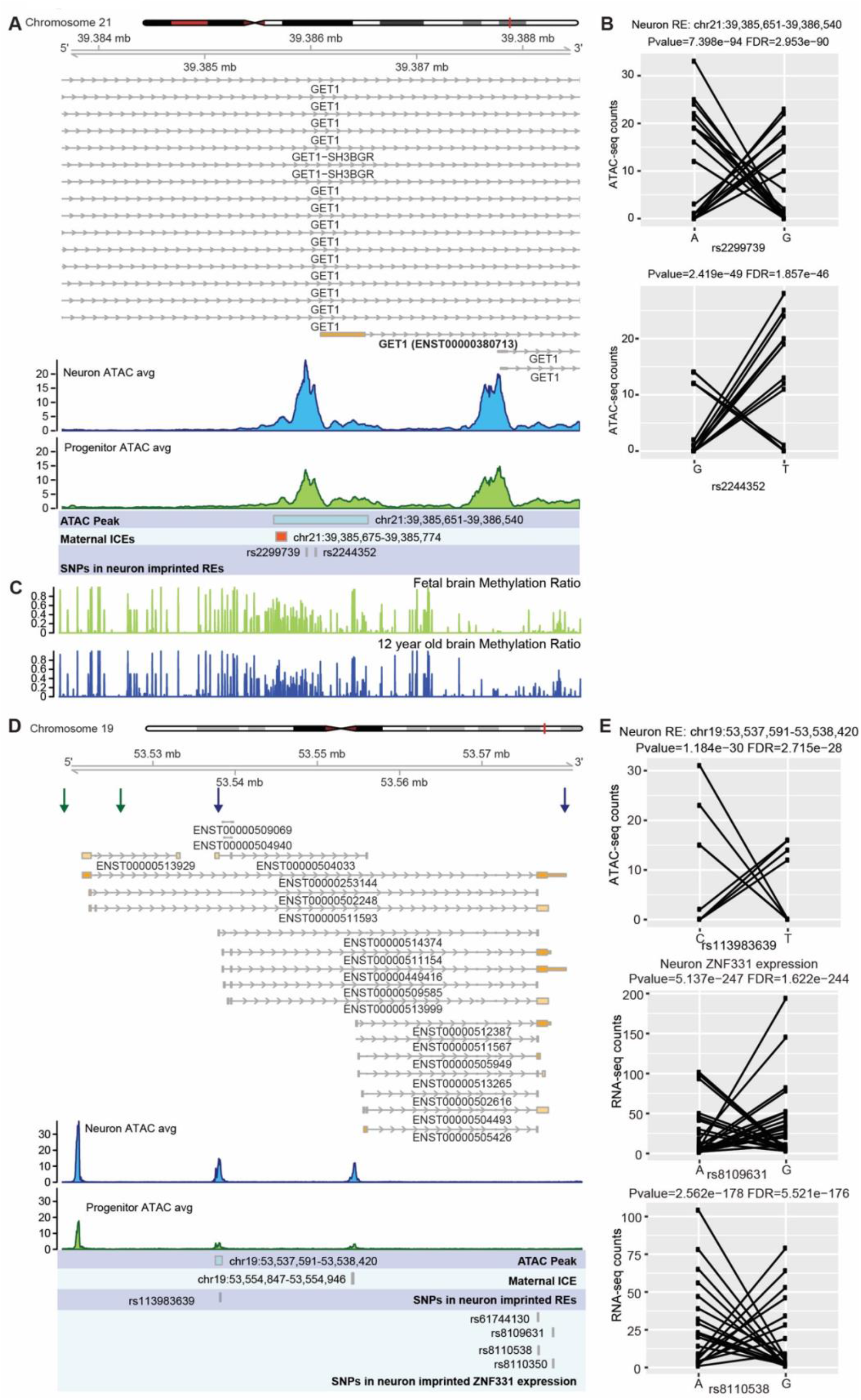
Imprinted REs and genes indicating isoform-specific imprinting in neurons. (A) Coverage plot of ATAC-seq in neurons and progenitors at GET1 locus. (B) Allelic ATAC-seq counts for the SNPs in the imprinted RE in (A). (C) Methylation ratio in human fetal brain and 12 year old brain (Lister et al. 2013). (D) Coverage plot of ATAC-seq in neurons and progenitors at the *ZNF331* locus. Boundaries of *ZNF331* isoforms showed neuron-specific imprinted expression patterns are indicated by blue arrows. Boundaries of *ZNF331* imprinted isoforms in LCL are indicated by red arrows. (C) Allelic ATAC/RNA-seq counts for SNPs in the imprinted RE and gene in (D).

We also found the gene *ZNF331* had cell-type and isoform-specific imprinted expression patterns (Figure 4D and 4E; Supplementary Figure 2D). We found a neuron-specific IRE (chr19:53,537,591-53,538,420) overlapped with the promoter of a subset of isoforms of *ZNF331* and these isoforms showed neuron-specific imprinted expression patterns (Figure 4E, boundaries of the imprinted isoforms are indicated by blue arrows). We also found a germline DMR near the promoters of the isoforms that could serve as an ICE of this region (Supplementary Figure 2C-D; Guo et al. 2014). The germline DMR is more methylated in oocyte and morula as compared to sperm (Supplementary Figure 2D), suggesting maternal imprinting. However, in progenitors, SNPs in this locus did not pass the threshold for significance and/or AR, so it was not possible to test for imprinted expression. Notably, isoform-specific imprinting of *ZNF331* was previously reported in multiple cell types and tissues, including the brain and LCLs (boundaries of LCL imprinted isoforms are indicated by red arrows in Figure 4D) (Court et al. 2014; Jadhav et al. 2019; Daelemans et al. 2010; Ben-David, Shohat, and Shifman 2014). These results indicate cell-type specific allelically biased isoform expression during neuronal development and differentiation.

### Progenitor-specific *DLK1* imprinting at Kagami Ogata/Temple syndrome paternal uniparental disomy locus

Uniparental disomy (UPD) results from homologous chromosomes, or parts of chromosomes, being inherited from only one parent (Robinson 2000). UPD and copy number variation mimicking UPD at imprinted sites results in abnormal expression. Copy number variation in the 14q32 imprinted gene cluster can lead to distinct maternal or paternal UPD phenotypes, named Temple syndrome and Kagami Ogata syndrome, respectively (Beygo et al. 2015; Buiting et al. 2008; Chen et al. 2005; Kagami et al. 2008). Individuals with Temple syndrome (UPD(14)mat) have characteristic features including pre- and postnatal growth retardation and developmental delay (Ioannides et al. 2014). Kagami Ogata syndrome (UPD(14)pat) results in prenatal overgrowth, developmental delay, and facial abnormalities with full cheeks and protruding philtrum (Ogata and Kagami 2016; Rosenfeld et al. 2015). Maternal deletions in the genomic region containing maternally expressed genes *MEG3, MEG8* and *RTL1* and sometimes containing paternally expressed gene *DLK1* have been identified in individuals with Kagami Ogata syndrome (Rosenfeld et al. 2015). Generally, individuals with maternal deletions in this region lack expression of the maternally expressed genes, but show overexpression of *DLK1* in blood and placenta (Ogata and Kagami 2016). In Temple syndrome, *DLK1* expression is lost due to the paternal deletion of this locus (Prasasya et al. 2020). We found evidence for imprinting of *DLK1* gene expression in progenitors but not neurons (Figure 5C and 5D; Supplemental Figure 2E).

**Figure 5.**
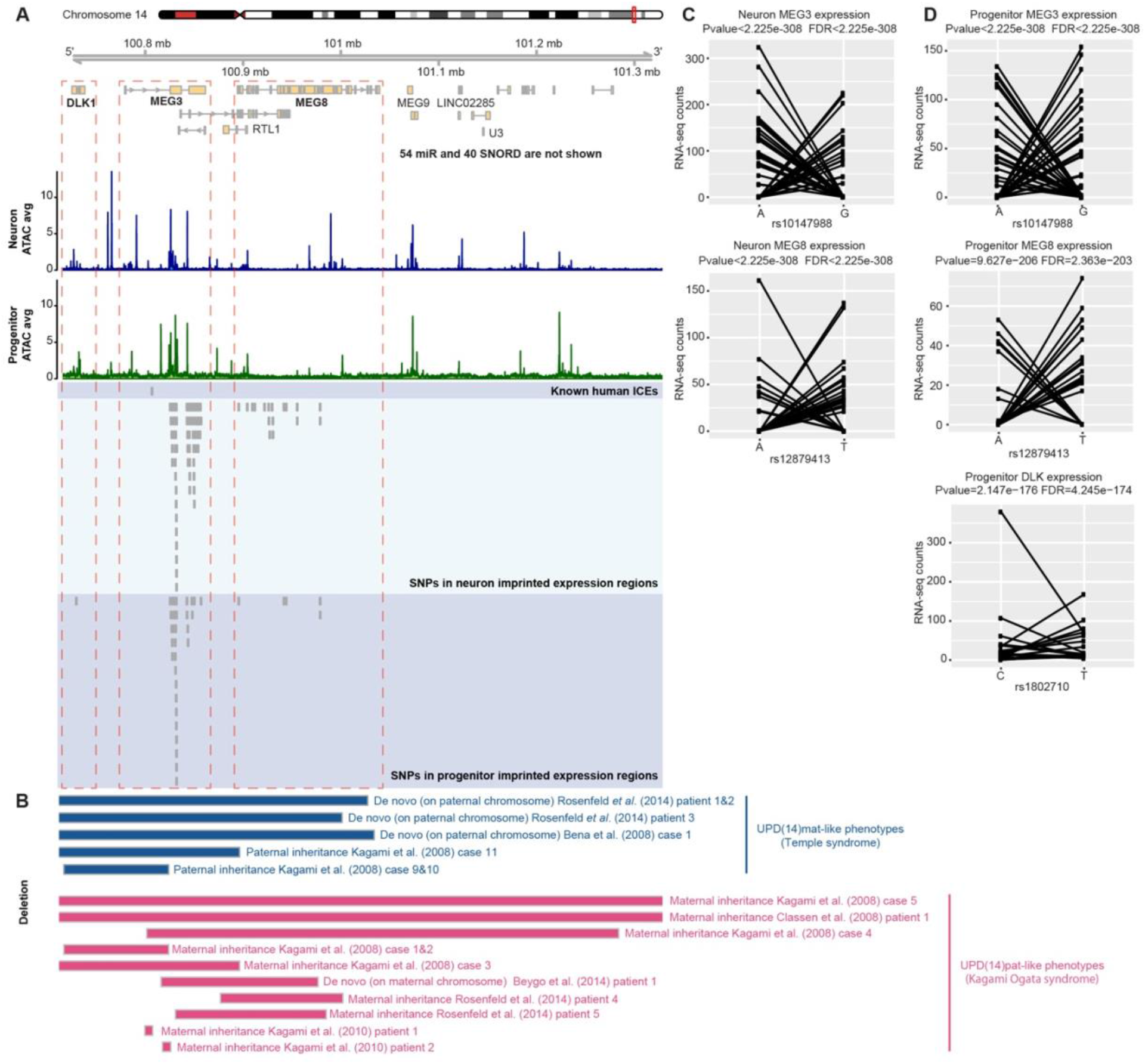
Progenitor-specific *DLK1* imprinting at Kagami Ogata syndrome paternal uniparental disomy locus. (A) Coverage plot of ATAC-seq in neurons and progenitors at Kagami Ogata syndrome paternal uniparental disomy locus. *DLK1* showed progenitor-specific imprinting expressions at genomic deletion regions. (B) Copy number variation in the 14q32 imprinted gene cluster related to Temple syndrome and Kagami Ogata syndrome. (C) Allelic RNA-seq counts for the SNPs in *MEG3* (rs10147988) and *MEG8* (rs12879413) in neurons. (D) Allelic RNA-seq counts for the SNPs in *MEG3* (rs10147988), *MEG8* (rs12879413), and *DLK1* in progenitors (rs1802710).

*DLK1*, previously known to be paternally expressed, promotes neurogenesis of neural progenitors in both mouse and human (Surmacz et al. 2012; Ferrón et al. 2011). In agreement with the function of *DLK1*, we found *DLK1* showed a significantly higher expression level in progenitors than in neurons (log2FC=-3.67, FDR = 6.5e-244). Here, we suggest that *DLK1* imprinting in progenitors contributes to the opposing phenotypes of overgrowth observed in Kagami Ogata syndrome, and growth retardation observed in Temple syndrome.

## DISCUSSION

Mutations at imprinted loci can lead to parent-of-origin dependent inheritance for neurodevelopmental disorders. In order to better explain the mechanism underlying parent-of-origin disorders, imprinting must be detected within relevant cell types. By combining high-throughput sequencing data (RNA-seq and ATAC-seq) with genotype data, we identified cell-type specific IREs and imprinted genes in two major cell types during human neuronal differentiation. We identified well-known IREs and genes in the PWS/AS region in neurons, providing confidence in our approach. We also identified cell-type specific IREs and genes as new candidates for genomic imprinting. We found cell-type specific REs may affect isoform-specific gene expression, as in neurons for *GET1*. Finally, we show that progenitor-specific imprinting of *DLK1* overlaps with deletions causing Kagami Ogata syndrome, suggesting neuronal progenitors contribute to neurobehavioral and growth changes observed in individuals with this syndrome.

Cell-type specific imprinted gene expression and chromatin accessibility showed dynamic changes of genomic imprinting during human neuronal differentiation. In addition to imprinted genes, we also identified IREs in neurons and progenitors, allowing us to explore the regulatory mechanisms of genomic imprinting. We found some genes only showed cell-type specific imprinted promoters but not imprinted expression, suggesting that the IREs are established prior to imprinted expression. We also found the IREs showed higher GC content and higher overlap with CpG islands, suggesting methylation of these REs is likely the regulatory mechanism underlying the imprinted signal in chromatin accessibility.

Although previous studies showed loss of imprinting in human pluripotent stem cells cultured *in vitro* (Bar et al. 2017; Frost et al. 2011), another study on induced pluripotent stem cells suggested genomic imprinting is not erased at the AS/PWS locus (Chamberlain et al. 2010). In this study, the well-known imprinted genes and promoters at AS/PWS locus were found in the neurons derived from neural progenitor cells in this study (Figure 2). The identification of these well-known imprinted genes indicated that the primary cell culture system used here was sufficient to study genomic imprinting at certain loci. However, it is unknown whether loss of imprinting at other sites occurred due to cell culture conditions.

Here, we inferred genomic imprinting using allelic ratios of RNA-seq and ATAC-seq reads. We separated genomic imprinting from allelic effects using a threshold for the number of individuals with extreme allelic ratio. A difficulty of identifying genomic imprinting using next generation sequencing data without genotype data from parents is to separate RMAE/RMACA from imprinted genes/REs. RMAE are mostly found in chromosome X due to the X-chromosome inactivity, and there are less than 5% of genes showing RMAE on autosomes (Kravitz and Gregg 2019). In this study, we removed the previously known RMAE/RMACA (Xu et al. 2017; Gimelbrant et al. 2007) to increase the confidence of imprinting calls. However, parental genotype data are necessary to completely disambiguate RMAE/RMACA from imprinting and so we refer to all novel imprinted genes/REs as putatively imprinted.

Additional experimental validation is also necessary to detect the molecular mechanisms of genomic imprinting during human neuronal differentiation. Genomic editing (deletion or modification) for IREs can be used to study their regulation of imprinting. Finally, we envision that combining the cell-type specific genomic imprinting with well-powered parent-of-origin genome-wide association studies (GWASs) or parent-of-origin rare variant association studies, will allow a better understanding of parent-of-origin effects on neurodevelopmental disorders.

## METHODS

### Cell culture of primary human neural progenitor cells (phNPCs)

We cultured and differentiated the phNPCs into neurons following the same methods in our previous work (Stein et al. 2014; Liang et al. 2021).

### ATAC-seq and RNA-seq library preparation for human neural progenitors and neurons

ATAC-seq libraries were prepared immediately following cellular dissociation described in our previous methods (Buenrostro et al. 2015; Liang et al. 2021). All libraries were sequenced on an Illumina HiSeq2500 or MiSeq machine using 50 bp paired-end sequencing. RNA-seq libraries were prepared as previously described (Aygün et al. 2021) and were sequenced on a NovaSeq S2 flow cell using 150 bp paired-end sequencing.

### ATAC-seq, RNA-seq, and genotype data pre-processing

Raw ATAC-seq and RNA-seq data were quality controlled and aligned to the human genome (GRCh38/hg38) using WASP to prevent mapping bias as previously described (Liang et al. 2021; Aygün et al. 2021). Genotype data were preprocessed and imputed as previously described (Liang et al. 2021).

### Allele-specific read counts

We used GATK tools (McKenna et al. 2010) to extract allele-specific read counts for every bi-allelic SNP (in accessible peaks or expression regions). We first filtered for SNPs within each donor that had sufficient read depth by retaining SNPs with total counts greater than or equal to 10 for neuron and progenitor samples, separately. Then to calculate allelic imbalance in chromatin accessibility and gene expression, we retained those SNPs with average read counts for all heterozygous donors greater than or equal to 15 for chromatin accessibility and 30 for gene expression. Finally, we retained only those SNPs that meet these previous thresholds for at least 5 heterozygous donors.

### Estimation of over-dispersion and identification of imprinted chromatin accessibility and gene expression

We identified over-dispersion using a likelihood ratio test based on the beta-binomial distribution for the allelic count (# reads from the reference allele given the # reads from the reference allele and alternative allele) at each SNPs. The allelic count can be modelled by a beta-binomial distribution, with the probability of expressing the parental-specific allele modelled by beta distribution (accounting for over-dispersion), and the number of reads observed modeled by a binomial distribution. The apeglm Bioconductor package was used to estimate parameters (Zitovsky and Love 2019). The likelihood ratio test was used to determine significance for the over-dispersion parameter for the heterozygous SNPs. To identify the imprinted chromatin accessibility and gene expression, we identify the SNPs in imprinted accessible or expression regions using the following three conditions: 1) SNPs have at least 2 donors with an allelic ratio greater than or equals to 0.8; 2) SNPs have at least 2 donors with an allelic ratio less than or equals to 0.2; and 3) SNPs have significant over-dispersion of the allelic ratio (FDR < 0.05 (Benjamini and Hochberg 1995)).

SNPs in the RMAE gene body and promoter regions were removed (Gimelbrant et al. 2007). For RMACA from mouse cells (Xu et al. 2017), we converted the RMACA regions from mouse genome (mm9) to the human genome (hg38) using liftOver from UCSCtools via the R package (rtracklayer v1.44.0).

### TFBS enrichment analysis

Potential transcription factor binding sites were called in the human genome using TFBSTools from the JASPAR2016 core database as previously (Liang et al. 2021). We calculated the enrichment of TFBS in Neuron/Progenitor IREs using the binomial test (McLean et al. 2010). First, we found accessible regions (n) overlapping with all TFBSs for a given TF and calculated the fraction of base pairs of the motif compared to the overall base pairs of accessible peaks (p). Then, we counted the number of IREs (k) overlapping with TFBSs for this TF (k). The final step was to calculate P=Pr_binom_(x>=k|n,p) using the binomial test to get the significance of the enrichment. We further filtered the enrichment results by differential expression from the same set of cells, and only kept the TFs with cell-type specific significantly enriched in imprinted REs and significantly differentially expressed in the cell type (Aygün et al. 2021).

### Identification of differential methylation regions among sperms, oocytes and morula

The genome-wide DNA methylation profiles for sperms (N=4), oocytes (N=2) and morula (N=3) were downloaded via NCBI Gene Expression Omnibus (GEO) under accession number GSE49828. R package ‘MethylKit’ (v1.16.1) was used to analyze differential methylation regions for sperms vs oocytes and morula or oocytes vs sperm and morula. After filtering the bases with < 5 reads, 100-bp-tiles was called and the methylation level was estimated as previously described (Akalin et al. 2012). A logistic regression model is used to identify the differential methylation tiles. The p values were adjusted to q values by the SLIM method (H.-Q. Wang, Tuominen, and Tsai 2011). Tiles with q-value<0.05 and percent methylation difference larger than 25% were assigned as differential methylation regions.

### DATA ACCESS (including public database accession numbers for all newly generated data and/or reviewer links to deposited data when accessions are not yet public. Previously published accessions should be included in the Methods section where appropriate)

ATAC-seq/RNA-seq data and genotype data for neurons and progenitors are available via dbGAP (ph001958 and phs2493).

#### Human CpG island

https://genome.ucsc.edu/cgi-bin/hgTables?hgsid=578954849_wF1QP81SIHdfr8b0kmZUOcsZcHYr&clade=mammal&org=Human&db=hg38&hgta_group=regulation&hgta_track=knownGene&hgta_table=0&hgta_regionType=genome&position=chr9%3A133252000-133280861&hgta_outputType=primaryTable&hgta_outFileName=

#### Genome-wide DNA methylation profiles for sperms, oocytes and morula

https://www.ncbi.nlm.nih.gov/geo/query/acc.cgi?acc=GSE49828

#### Imprinted genes

https://geneimprint.com/site/genes-by-species

#### Genome-wide DNA methylation profiles for brain tissues

https://www.ncbi.nlm.nih.gov/geo/query/acc.cgi?acc=GSE47966

## ACKNOWLEDGMENTS

This work was supported by NIH (R00MH102357, U54EB020403, R01MH118349, R01MH120125), and Brain Research Foundation to JLS.

## Author Contributions

JLS and MIL conceived the study. JLS directed and supervised the study. JLS provided funding. NA performed pre-processing of allele specific gene expression data. MIL assisted with statistical identification of imprinting REs and genes. FI assisted with interpretation of the results. DL performed pre-processing of ATAC-seq data and identification of imprinting REs and genes. JLS and DL wrote the manuscript. All authors commented on and approved the final version of the manuscript.

## DISCLOSURE DECLARATION (including any conflicts of interest)

The authors do not declare any conflicts of interest.

## Supplementary Tables

**Table S1**. Imprinted REs in neurons and progenitors. VariantID is the name of the caSNP. RefAllele and altAllele are reference allele and alternative allele of the caSNP. Theta.hat is the estimate of dispersion using the apeglm Bioconductor package’s bbEstDisp function (larger theta indicating less variance), p.hat and p.hat0 are the fitted allelic ratio for the full and reduced models respectively, and stat is the likelihood ratio test statistic. P.Value is the nominal p-value for the test of overdispersion; adj.P.Val is the Benjamini-Hochberg FDR adjusted p-value. H_Het_donor is the number of heterozygous donors for the SNP. counts are the total counts for the SNPs across all donors. Chr, peakstart and peakend are the coordinates (hg38) for the REs. Promotor represents the overlapped promoters with the RE. KnownImp if true means the RE overlaps with the promoter of known imprinted genes. The NeuronImprintedREs tab is for nIREs and the ProgenitorImprintedREs tab is for pIREs.

**Table S2**. Imprinted genes in neurons and progenitors. SNPIDs is the name of the caSNP. RefAllele and altAllele are reference allele and alternative allele of the caSNP. Theta.hat, p.hat, p.hat0 and stats are as defined for Supplementary Table 1. P.Value is the nominal p-value of dispersion; adj.P.Val is the Benjamini-Hochberg FDR adjusted p-value. H_Het_donor is the number of heterozygous donors for the SNP. counts are the total counts for the SNPs across all donors. Gene is the gene name. KnownImp if true means the gene is a known imprinted gene. The NeuronImprintedGenes tab is for neuron imprinted genes and the ProgenitorImprintedGenes tab is for progenitor imprinted genes.

---

**Supplemental Figure 1.**
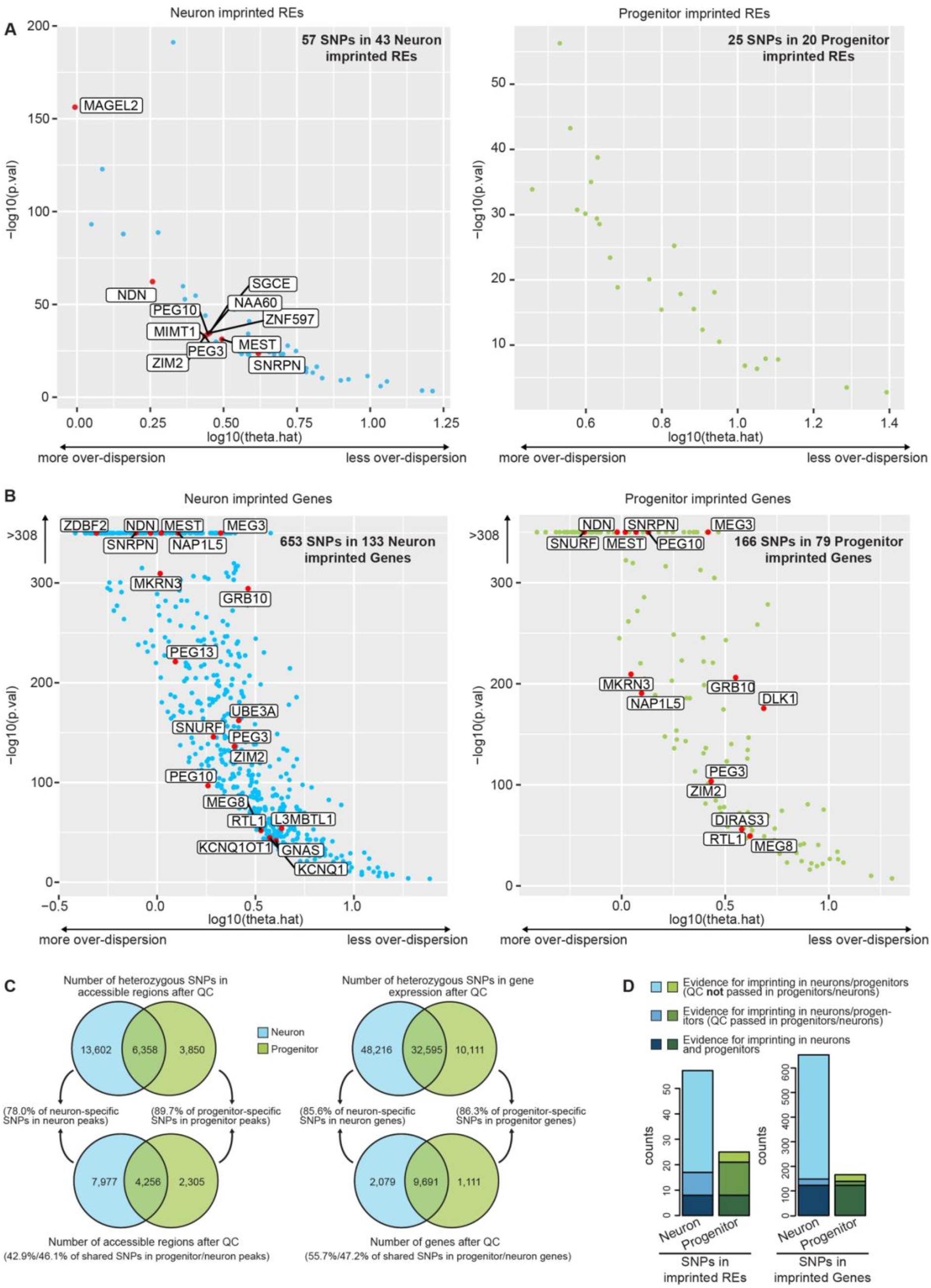
(A) Dispersion level and the p values for SNPs in IREs. IREs overlapping with the promoters of known imprinted genes are labeled by red dots. (B) Dispersion level and the p values for SNPs in imprinted genes. (C) Comparison of SNPs in gene expression region and REs (before and after QC) between neurons and progenitors. (D) Comparison of SNPs in imprinted gene expression region and REs between neurons and progenitors.

**Supplemental Figure 2.**
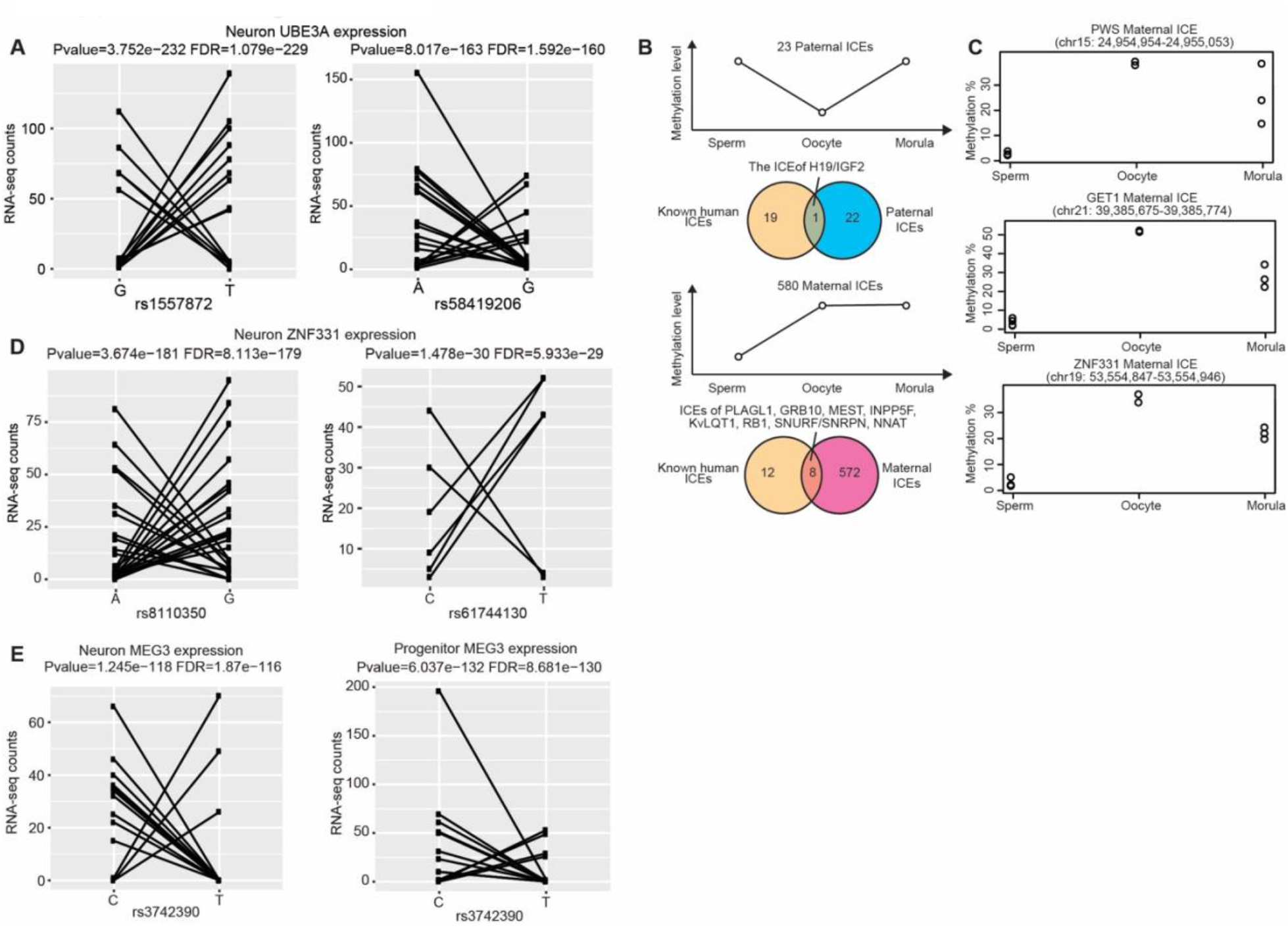
(A) Allelic ATAC-seq counts for the SNPs in the expression region of *UBE3A*. (B) Schematic cartoon of paternal (23) and maternal (580) ICEs and overlaps of ICEs and promoters of known imprinted genes. (C) Methylation rates of ICEs in sperm, oocytes and morula. (D) Allelic ATAC-seq counts for the SNPs in the expression region of *ZNF331*. (E) Allelic ATAC-seq counts for the SNPs in the expression region of *MEG3*.

## Notes

### Competing Interest Statement

The authors have declared no competing interest.

### Summary of Updates

Uploaded supplemental tables.

